# Bivariate analysis of barley scald resistance with relative maturity reveals a new major QTL on chromosome 3H

**DOI:** 10.1101/748368

**Authors:** Xuechen Zhang, Ben Ovenden, Beverley A. Orchard, Meixue Zhou, Robert F. Park, Davinder Singh, Andrew Milgate

## Abstract

The disease scald of barley is caused by the pathogen *Rhynchosporium commune* and can cause up to 30-40% yield loss in susceptible varieties. In this study, the Australian barley cultivar Yerong was demonstrated to have resistance that differed from Turk (*Rrs1*) based on seedling tests with 11 *R. commune* isolates. A doubled haploid population with 177 lines derived from a cross between Yerong and Franklin was used to identify quantitative trait loci (QTL) for scald resistance. Scald resistance against four pathogen isolates was assessed at the seedling growth stage in a glasshouse experiment and at the adult growth stage in field experiments with natural infection over three consecutive years. A QTL on chromosome 3H was identified with large effect, consistent with a major gene conferring scald resistance at the seedling stage. Under field conditions, scald percentage was negatively correlated with early relative maturity. A bivariate analysis was used to model scald percentage and relative maturity together, residuals from the regression of scald percentage on relative maturity were used as our phenotype for QTL analysis. This analysis identified one major QTL on chromosome 3H, which mapped to the same position as the QTL identified for scald resistance at seedling stage. The identified QTL on 3H is proposed to be different from the *Rrs1* on the basis of seedling resistance against different *R. commune* isolates and physical map position. The analysis also identified an additional novel QTL on chromosome 7H. This study increases the current understanding of scald resistance and identifies genetic material possessing QTLs useful for the marker-assisted selection of scald resistance in barley breeding programs.

## Introduction

Scald is a serious foliar disease in barley (*Hordeum vulgare*) that is caused by *Rhynchosporium commune*. The pathogen can cause up to 30-40% yield loss in susceptible varieties and is found in all barley-growing regions worldwide ^1^. Control of scald disease requires a multi-facetted approach, including application of fungicides, cultural disease management, manipulation of sowing date and the use of resistant cultivars ^2^. *R. commune* populations have changed rapidly in response to newly-developed fungicides and resistant plant varieties ^3–6^. One of the most sustainable strategies for *R. commune* management is to develop and deploy disease-resistant barley cultivars through the introgression and pyramiding of different resistance genes (major or minor). Traditional methods of phenotypic selection for complex patho-systems such as scald can be improved through detailed genetic studies, which allow the implementation of marker-assisted selection (MAS) in breeding programs.

Scald resistance is governed by both major and minor genes. Major resistance genes provide high levels of resistance at all plant growth stages, while minor resistance genes generally provide partial levels of resistance at the adult plant stage ^7,8^. Reduced scald symptoms in adult stage plants under field conditions might also result from disease escape through physical barriers to infection ^9^. In terms of scald resistance in barley, flowering time, plant height and canopy structure can affect scald symptoms by physically limiting the upward spread of the splash-dispersed pathogen ^8^. The major scald resistance genes discovered so far have been mainly identified through experiments with seedlings via inoculation with specific isolates ^8^. The problem with using major scald resistance genes in breeding programs is a lack of durability. Quantitative genes are thought to be more durable, and it has been suggested that pyramiding these genes could reduce the ability of *R. communes* to rapidly acquire new virulence combinations ^10^.

A large number of QTLs for scald resistance have been discovered in barley. Two genomic regions in particular have been frequently associated with resistance; the *Rrs1* locus on chromosome 3H and the *Rrs2* locus on chromosome 7H ^8,11^. These two loci have been detected in different mapping populations, across different environments, under glasshouse conditions after inoculation with specific isolates, and under field conditions ^8,11^. Progress of our understanding of these loci has been made through fine mapping studies ^12,13^. It remains unknown if the QTLs detected at each of these two loci are alleles of the same gene, or if they are part of a closely linked gene cluster at each locus ^14,15^.

For the *Rrs1* locus, multiple major and minor scald resistance genes or QTLs have been identified, supporting its importance in barley germplasm worldwide ^12,16–21^. *Rrs1* was the first scald resistance gene reported in barley ^22^ with an associated RFLP marker of cMWG680, positioned at 455.3 Mb on chromosome 3H on the barley physical map ^18^. The location of the *Rrs1* locus was further fine mapped including all known markers to an interval of less than 9 Mbp at 448.4 Mb from Spanish barley varieties ^12^.

In this study, we performed screening against 11 *R. commune* isolates at seedling stage to demonstrate that resistance in the variety Turk (international source of *Rrs1*) differs from that resistance in the variety Yerong. These results contradicted previous findings that suggested Yerong carries a scald resistance QTL at the *Rrs1* locus on chromosome 3H ^23^. To further investigate this finding, screening of a Yerong/Franklin doubled haploid population was conducted for seedling resistance against four *R. commune* isolates and adult plant resistance under natural field conditions across three years. To overcome potential confounding effects of differences in the onset of flowering (determined as relative maturity) on scald resistance QTL detection within the population, a linear mixed model with a bivariate approach was used to analyse the field data. Using this approach, a major QTL on chromosome 3H from Yerong was detected for scald resistance at both seedling and adult plant stage. The QTL detected did not map to the *Rrs1* locus, suggesting the presence of a new and useful scald resistance QTL in Yerong.

## Materials and Methods

### Differential varieties screening

The seedling scald resistance of different barley varieties was tested against 11 *R. commune* isolates from the Wagga Wagga Agricultural Institute fungal isolate collection. These isolates were collected from southern New South Wales between 2013 and 2016 (Table 1). The *R. commune* isolates were grown on lima bean agar (LBA) at 20°C under 24 hour light condition ^7^. After 2 to 3 weeks, spores were harvested with a scalpel blade and the spore solutions for spray inoculation were adjusted to 2×10^6^ spores ml^−1^ in distilled water using a haemocytometer. Seeds of varieties were sown in 6-cm pots arrayed in a randomized complete block design consisting of 4 columns by 30 rows with three replicates of each genotype in each experiment for a total of 120 pots. A total of 40 varieties were included in the differential varieties screening, however, only the results for six key varieties (Yerong, Franklin, Turk, Atlas, Atlas46 and Litmus) were reported in this study (Table 2). Each variety by isolate combination was evaluated in at least two experiments. The responses of Turk (*Rrs1*), Atlas (*Rrs2*) and Atlas46 (both *Rrs1* and *Rrs2*) were compared with that of Yerong, while Litmus was used as the susceptible check variety. The barley varieties were uniformly sprayed with spore solutions at the 3-leaf-stage, and kept in a dark chamber for 48 hours at 18°C with 100% humidity. The seedlings were scored 14 days post-inoculation with a scoring scale from 1 to 5 (1 = resistant and 5 = very susceptible) following the methods of Jackson and Webster ^24^.

**Table 1.** Origin of isolates of scald from the Wagga Wagga Agricultural Institute fungal isolate collection used for screening

| Isolate | Host variety | Location | Collection date |
| --- | --- | --- | --- |
| WAI453 | Franklin | Wagga Wagga, NSW | 2013 |
| WAI1245 | Franklin | Wagga Wagga, NSW | 2013 |
| WAI2439 | Unknown | Downside, NSW | 2015 |
| WAI2463 | Buloke | Bogan Gate, NSW | 2015 |
| WAI2464 | Buloke | Bogan Gate, NSW | 2015 |
| WAI2466 | Buloke | Bogan Gate, NSW | 2015 |
| WAI2470 | Unknown | Mirrool, NSW | 2015 |
| WAI2471 | Unknown | Mirrool, NSW | 2015 |
| WAI2473 | Unknown | Mirrool, NSW | 2015 |
| WAI2636 | Buloke | Bogan Gate, NSW | 2015 |
| WAI2840 | Unknown | Finley, NSW | 2016 |

**Table 2.** Predicted values (BLUPs) with 95% confidence intervals for seedling resistance of different barley varieties against 11 different *R. commune* isolates

| Variety | Resistance | WAI453 | WAI1245 | WAI2439 | WAI2463 | WAI2464 | WAI2466 | WAI2470 | WAI2471 | WAI2473 | WAI2636 | WAI2840 |
| --- | --- | --- | --- | --- | --- | --- | --- | --- | --- | --- | --- | --- |
| Atlas | *Rrs2* | 2.4 ± 0.8 | 3.3 ± 0.9 | 2.3 ± 0.8 | 3.1 ± 0.8 | 4.0 ± 0.9 | 2.4 ± 0.6 | 2.8 ± 0.8 | 2.2 ± 0.8 | 2.9 ± 1.2 | 2.5 ± 0.5 | 3.5 ± 0.9 |
| Atlas46 | *Rrs1+Rrs2* | 2.1 ± 0.8 | 2.0 ± 0.8 | 2.3 ± 0.8 | 2.2 ± 0.8 | 2.1 ± 0.9 | 1.9 ± 0.6 | 2.1 ± 0.9 | 2.0 ± 0.8 | 2.0 ± 0.8 | 1.8 ± 0.5 | 3.5 ± 0.8 |
| Franklin |  | 4.2 ± 0.8 | 4.3 ± 0.8 | 3.5 ± 0.8 | 4.4 ± 0.8 | 4.4 ± 0.8 | 4.7 ± 0.5 | 4.4 ± 0.8 | 3.4 ± 1.3 | 3.6 ± 0.8 | 4.3 ± 0.5 | 4.4 ± 0.8 |
| Litmus | Susceptible | 4.5 ± 0.8 | 4.4 ± 0.8 | 4.4 ± 0.9 | 4.5 ± 0.8 | 4.5 ± 0.8 | 5.2 ± 0.5 | 4.4 ± 0.8 | 4.0 ± 0.8 | 4.3 ± 0.8 | 4.1 ± 0.5 | 3.9 ± 1.2 |
| Turk | *Rrs1* | 2.0 ± 0.8 | 2.2 ± 0.8 | 3.8 ± 0.8 | 2.4 ± 0.8 | 2.0 ± 0.8 | 1.7 ± 0.5 | 2.0 ± 0.8 | 2.0 ± 0.8 | 2.0 ± 0.8 | 1.7 ± 0.5 | 4.1 ± 0.8 |
| Yerong |  | 3.1 ± 0.8 | 3.4 ± 0.8 | 3.2 ± 0.8 | 4.3 ± 0.8 | 4.3 ± 0.8 | 4.7 ± 0.5 | 4.3 ± 0.8 | 2.3 ± 0.8 | 3.2 ± 0.8 | 3.8 ± 0.5 | 4.3 ± 0.8 |

### Plant material for QTL analysis

A doubled haploid (DH) population of 177 lines from a cross between varieties Yerong and Franklin was used in this study. Seeds for each genotype were obtained from the University of Tasmania. Franklin is an Australian two-rowed malting quality variety, and Yerong is an Australian six-rowed feed quality variety. The genetic linkage map of the population comprised 28 microsatellites and 196 diversity arrays technology (DArT) markers assembled by Li and Zhou ^23^.

### Evaluation of seedling resistance to scald

Four different *R. commune* isolates, WAI2466, WAI2470, WAI2471 and WAI2473, were used in the seedling resistance screening in the glasshouse experiments (Table 1). All four isolates were collected from southern New South Wales and are held in the Wagga Wagga Agricultural Institute fungal isolate collection (Table 1). Inoculum preparation and inoculations were as outlined for the differential screening. Spores were diluted to 1×10^6^ spores ml^−1^ in distilled water using a haemocytometer. Each of the four isolates was tested in a separate experiment. For each experiment, the 177 DH lines as well as the parental varieties Franklin and Yerong and an additional 21 check varieties (for a total of 200 genotypes) were sown in 6-cm pots arrayed in a 24 column by 25 row randomized complete block design with three replicates of each genotype for a total of 600 pots. Each variety by isolate combination was tested with three different experiments with three technical replicates per experiment. Seedlings were scored 14 days post-inoculation with a scoring system from 1 to 4 (1 = resistant and 4 = very susceptible) following the methods of Wallwork and Grcic ^7^.

### Field screening

Field screening for scald resistance was conducted at Wagga Wagga Agricultural Institute (Wagga Wagga, New South Wales) in 2015, 2016 and 2017. All field trials were sown in May with a randomised complete block design with two replicates for each genotype. Each genotype was sown in 1.2 m rows with 0.4 m spacing between each row. The primary inoculum for *R. commune* infection was residual barley crop debris from the previous harvest. Overhead irrigation was used regularly to supplement rainfall throughout the growing season to enhance the development of disease. Experiments were subject to a strict weed control and crop nutrition regime to maximize yield potential. Assessment of disease was based on leaf symptoms using the percentage of infected leaf area ^6^. Relative maturity at the time of disease assessment was determined using the Zadoks decimal score for plant development ^25^. Final plant height was also measured at physiological maturity in the 2017 experiment.

### Statistical analysis

All data was analysed using the software package ASReml-R version 3 ^26^ in the R environment ^27^. A linear mixed model following the approach of Gilmour, et al. ^28^ was used to analyse the data for the differential variety screening experiments as follows (Model 1):

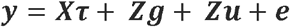

where ***y*** is the *n* × 1 vector of the response variable (scald score) across *p* = 22 experiments with each of the 11 *R. commune* isolates tested in a separate experiment, and that experiment repeated once. *n* = 2640 for the differential variety screening experiments as only selected varieties were included. ***τ*** is a *t* × 1 vector of fixed effects, including the overall mean scald score, corresponding to the *n* × *t* design matrix ***X***. The term ***g*** is the vector of genotypic random effects with associated design matrix ***Z*** used to model the genotype by experiment effects. The term ***u*** is the vector of random effects corresponding to the experimental design matrix ***Z***, which contains experiment-specific terms to capture extraneous variation including the experiment level blocking structure including replicate, row and column. The *n* × 1 residual vector ***e*** was modelled for each experiment.

A model similar to Model 1 above was also used to model scald scores for the seedling inoculation experiments with *n* = 7200 for the *p* = 12 experiments conducted.

For the field screening experiments measuring scald resistance and relative maturity, each of the three field experiments was modelled separately using a bivariate approach as follows:

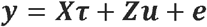

where ***y*** is a vector of length *n* = 2 × 479 containing stacked vectors for the two traits, *S*: scald resistance and *R*: relative maturity). ***τ*** is a vector of fixed effects including trait means and the trait by genotype effects for the design matrix ***X***. The term ***u*** is the vector of replicate, column and row effects for each trait corresponding to the experimental design structure ***Z***. The vector ***e*** of length *n* containing the residuals of the two traits *S* and *R* was modelled with a separable autoregressive process of order one (*AR*1 ⊗ *AR*1) and an unstructured variance-covariance matrix between traits. This structure permits the fitting of a linear relationship at the residual level between the two traits. In all models, the significance of fixed effects was assessed using the techniques of Kenward and Roger ^29^ and the significance of random effects other than ‘replicate’ was determined using log-likelihood ratio tests ^30^.

The linear relationship between scald resistance and relative maturity was determined from the trait:genotype covariance modelling after the general approach of Van Beuningen and Kohli ^31^ and the paired case-control study example detailed in Butler, et al. ^26^ as follows:

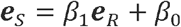

where the slope of the regression is calculated as:

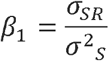

and the intercept *β*_0_ was determined from the overall BLUPs for the two traits *S* and *R* from the bivariate mixed model. The difference (residual) between the BLUP from the mixed model for scald resistance of a genotype and the predicted value on the trend line was calculated. These values are referred to hereafter as the deviation from the regression of scald resistance on relative maturity (DRSRRM).

### QTL analysis and positioning of identified QTL on the barley physical map

The DRSRRM values for each individual field experiment were used as the phenotype to detect QTL for scald resistance adjusted for the relationship between scald resistance and relative maturity. For the phenotype analyses of scald percentage, relative maturity and plant height BLUPs were obtained from the model for each experiment. Phenotypes were used for QTL analysis using the MapQTL6.0 software package ^32^. QTLs were first analysed with interval mapping (IM). The closest marker to each QTL was selected as a cofactor for multiple QTL mapping (MQM). A logarithm of the odds (LOD) threshold value of 3 was used to identify QTL.

The primer sequences of markers associated with the identified QTL for scald resistance and fine mapped *Rrs1* ^12^ were used to do BLAST searches by using the IPK Barley BLAST Server (http://webblast.ipk-gatersleben.de/barley). The barley pseudomolecules Morex V 2.0 2019, was used for the BLASTn search. Default settings were used to do the BLASTn search and the best hit was used to decide the physical position of the detected QTL.

## Results

### Differential varieties screening

Atlas46 (*Rrs1* + *Rrs2*) was resistant to all the isolates used in this study, with scores under 2.3, except isolate WAI2840, which was virulent on all varieties with a score of more than 3.5 (Table 2). Turk (*Rrs1*) was resistant to all the isolates used in this study with scores under 2.4, except two isolates WAI2439 and WAI2840. Varieties carrying *Rrs2* (Atlas and Atlas46) were resistant against isolate WAI2439 (Table 2). Varieties conferring *Rrs1* (Turk and Atlas46) showed higher level of resistance than Atlas and Yerong against two isolates, WAI1245 and WAI2464. Varieties carrying either *Rrs1* or *Rrs2* were resistant against isolates WAI453, WAI2466, WAI2471 and WAI2636.

Turk was significantly more resistant than Yerong against five isolates and equivalent against the remaining six isolates tested. Together, these results suggested that *Rrs1* from Turk was not present in Yerong. Compared to Yerong, Atlas showed a higher level of resistance against isolates WAI2439, WAI2463, WAI2466, WAI2470 and WAI2636, suggesting *Rrs2* is absent in Yerong. Yerong was resistant against WAI2471, and displayed moderate resistance against isolates WAI453, WAI1245, WAI2439 and WAI2473 with the scores between 3.0 and 3.4. Yerong showed a better resistance than Franklin against isolates WAI453, WAI1245 and WAI2471with the overall scores of Yerong being lower than those of Franklin. Litmus was susceptible to all the isolates in this study.

### QTLs for scald resistance at the seedling stage

There were no significant differences (using 95% confidence intervals) in resistance to the four isolates WAI2466, WAI2470, WAI2471 and WAI2473 among the two parent cultivars in the seedling stage screening experiments. However, there was phenotypic variation in resistance to the different isolates among the DH population lines (Fig. 1). The disease scores among DH population lines against WAI2466 showed a bimodal distribution of scores between resistant and susceptible lines. The variation of disease scores against WAI2470 was the lowest among all four isolates. The distribution of disease scores against WAI2471 was skewed towards higher levels of disease. In contrast, the distribution of disease scores against WAI2473 was skewed towards the lower end of the scoring scale.

**Fig. 1.**
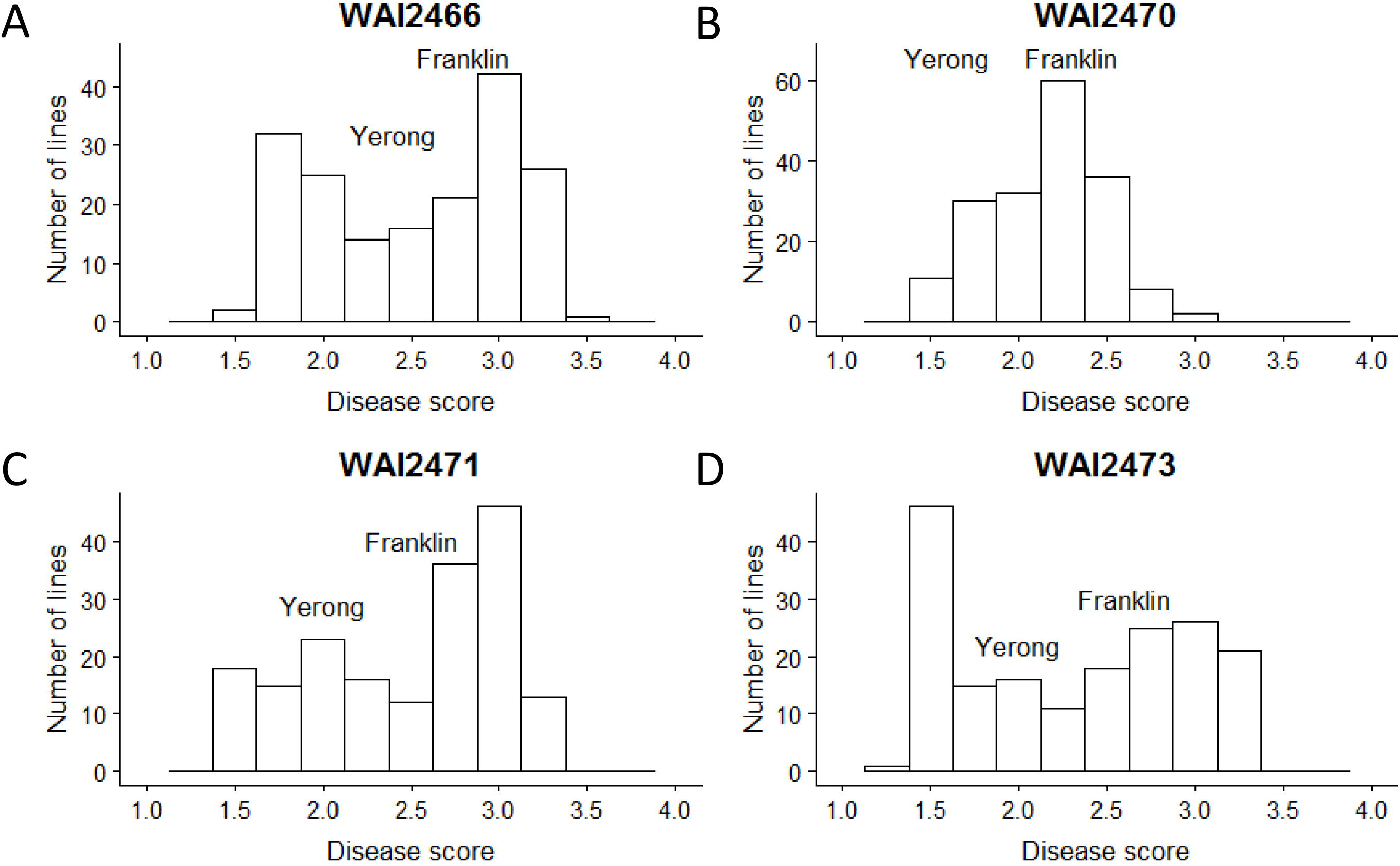
Frequency distribution of scald resistance to four pathogen isolates in the Yerong/Franklin population at the seedling stage different isolates. The positions of text labels “Yerong” and “Franklin” in the figure are based on the disease score of each parent at the seedling stage

One major QTL on chromosome 3H, designated as QTL-WAIYerong-3H, was identified for seedling resistance to all four different *R. commune* isolates (WAI2466, WAI2470, WAI2471 and WAI2473) at the same position (Table 3) at 178.7 Mb. Flanking markers of this QTL, bPb-0068 and Bmag0006, are mapped at 120.3 and 178.7 Mb on the physical map (Fig. 2). The LOD scores varied from 23.8 to 38.7, explaining more than 46% of the phenotypic variation (Table 3).

**Table 3.**
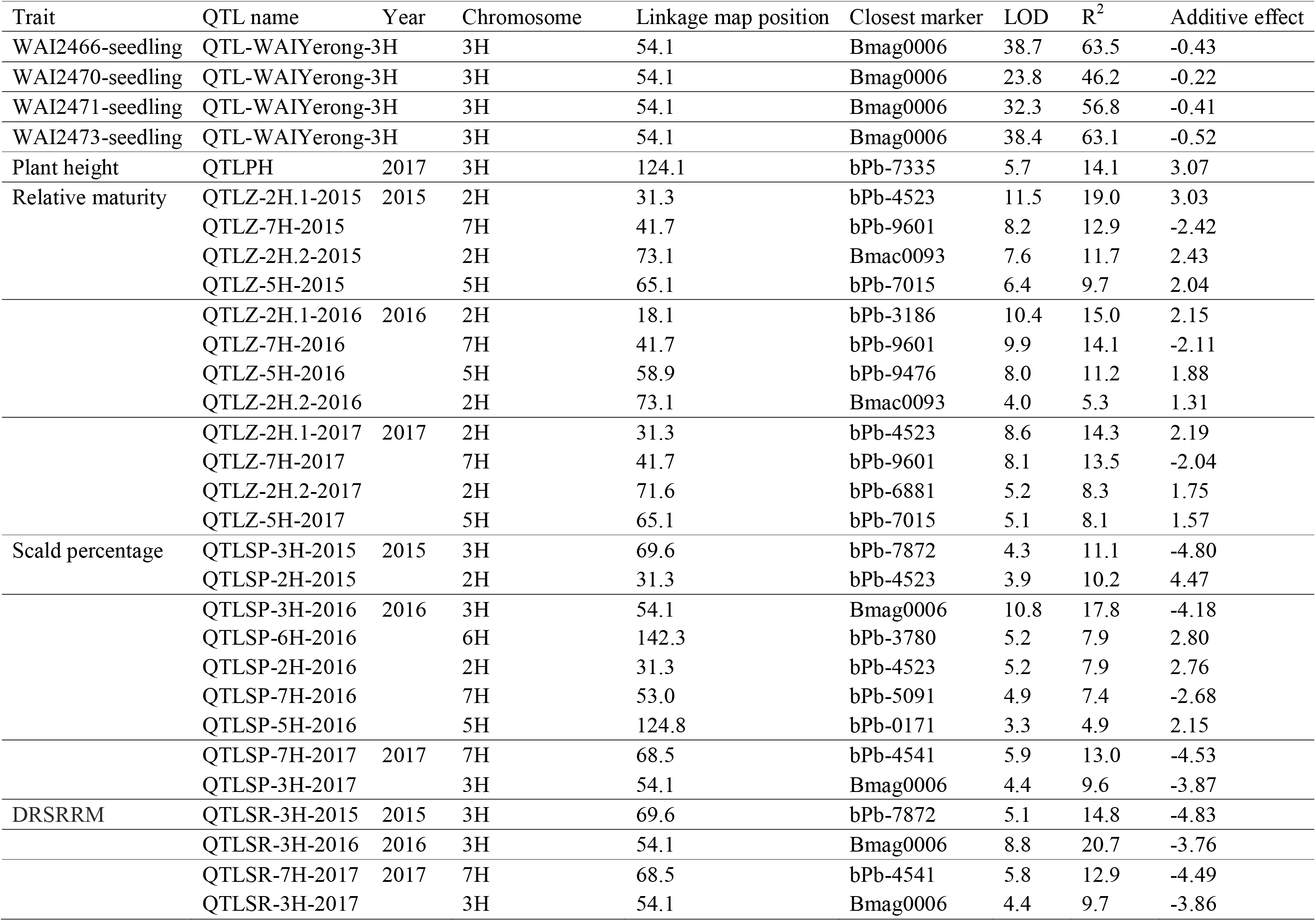
Summary of QTLs for scald resistance at the seedling stage to four different isolates: WAI2466, WAI2470, WAI2471 and WAI2473, and Summary of the QTLs for all traits measured in the field experiments

**Fig. 2.**
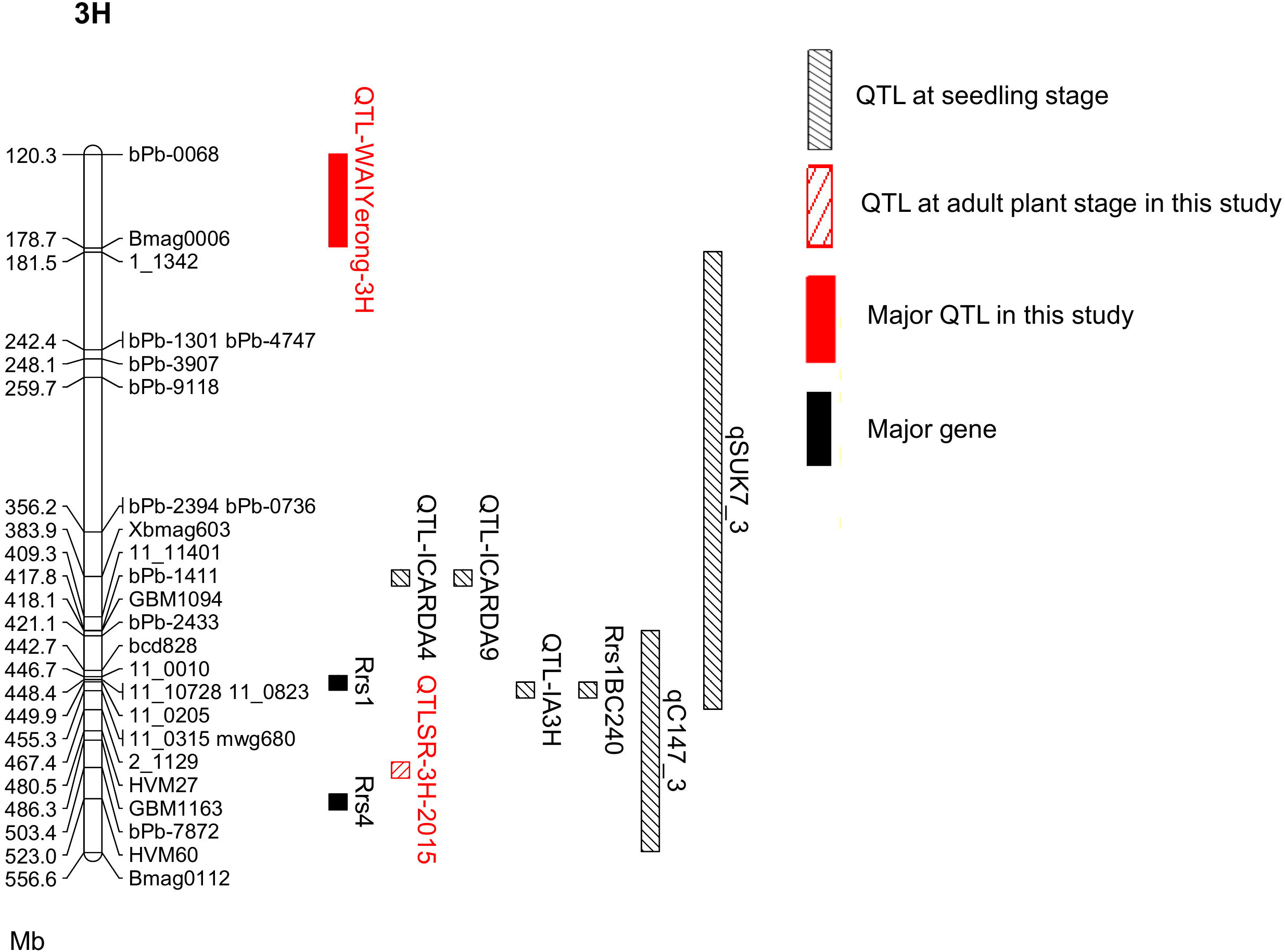
Positions of marker sequences on 3H pseudomolecule (100 - 600 Mb) of Morex genome assemble version 4. Flanking markers of the major QTL on chromosome 3H from Yerong identified in this study (bPb-0068 and Bmag0006) mapped at 120.3 and 178.7 Mb on the physical map. Flanking markers of QTLSR-3H-2015, bPb-7872 and bPb-8410 mapped at 486.2 and 503.4 Mb on the physical map. The physical position of fine mapped *Rrs1* is identified using flanking markers, at 448.4 Mb ^12^. The original RFLP marker mwg680, which is closely linked to the *Rrs1* gene, is located at 455.3 Mb ^50^. The resistance QTLs from ICARDA4 and ICARDA9 are mapped at 383.9 Mb based on marker Xbmag603 ^17^. The resistance QTL Rrs1BC240from wild barley H. *spontaneum* CPI 109853 is mapped at 455.3 Mb ^17^. The locations of resistance QTLs identified from Steptoe (qSUK7_3) and CIho 3515 (qC147_3) are cited from Coulter, et al. ^20^. The resistance QTL QTL-IA3H from Abyssinian is mapped at 455.3 Mb against four isolates ^19^. Major resistance gene *Rrs4* is mapped at 523.0 Mb based on marker HVM60 ^51^

### Scald resistance under field conditions

The parent varieties Franklin and Yerong showed similar levels of scald resistance and relative maturity across the different years (Table 4). There were no significant differences between Franklin and Yerong for any of the traits measured across the different years (p > 0.01). However, the DH lines showed substantial phenotypic variation in scald resistance and relative maturity (Fig. 3). Average disease incidence was more severe in 2016, reaching 43.0%, higher than the average scald percentages in 2015 (29.0%) and 2017 (22.1%) (Table 4).

**Table 4.** Phenotypic predicted values (BLUPs) of traits measured in the Yerong/Franklin population under field conditions

| Trait | Year | Average | Max | Min | Franklin | Yerong |
| --- | --- | --- | --- | --- | --- | --- |
| Relative maturity | 2015 | 42.1 | 58.4 | 33.3 | 38.2 | 38.3 |
|  | 2016 | 47.0 | 59.5 | 37.0 | 42.8 | 45.9 |
|  | 2017 | 45.2 | 61.0 | 35.5 | 43.0 | 41.5 |
| Scald percentage | 2015 | 29.0 | 63.4 | -0.3 | 23.7 | 21.7 |
|  | 2016 | 43.0 | 63.5 | 15.1 | 31.6 | 34.3 |
|  | 2017 | 22.1 | 61.5 | 6.5 | 16.0 | 18.5 |
| DRSRRM | 2015 | -0.1 | 34.8 | -26.4 | -2.2 | -4.3 |
|  | 2016 | -0.2 | 19.7 | -18.6 | -7.6 | -7.8 |
|  | 2017 | -0.1 | 39.3 | -15.6 | -6.1 | -3.6 |
| Plant height | 2017 | 96.3 | 115.5 | 72.0 | 107.0 | 106.0 |

**Fig. 3.**
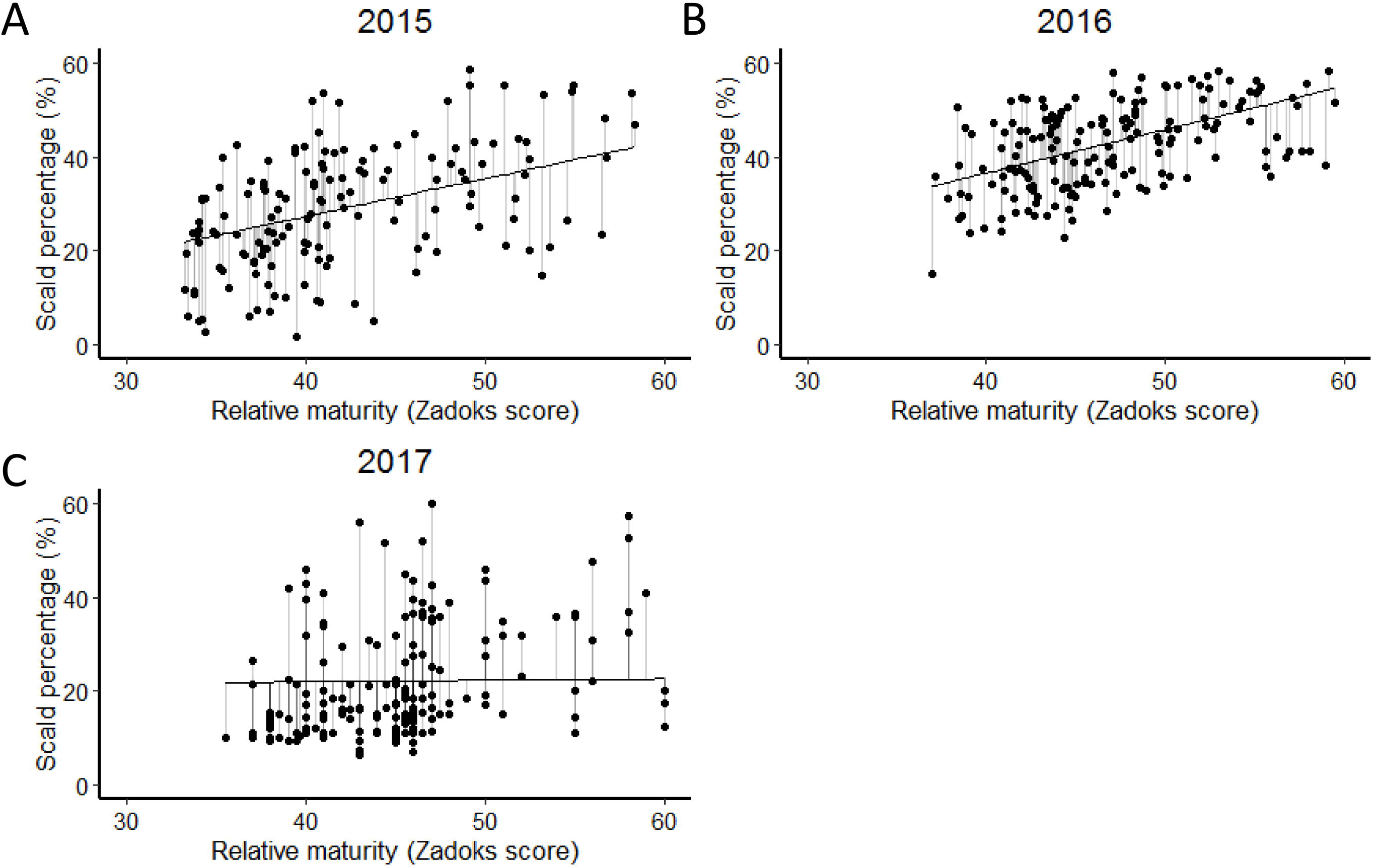
The scald percentage in the DH population of Yerong/Franklin plotted against Zadoks score

A significant correlation (Pearson’s correlation coefficient) between scald percentage and relative maturity was observed among the DH lines (Fig. 3) in 2015 (r = 0.50, p < 0.01), 2016 (r = 0.51, p < 0.01). Later flowering lines (with lower relative maturity scores) tended to have a lower scald percentage. In 2017, although Pearson correlation coefficient between relative maturity and scald percentage was significant (r = 0.34, P < 0.01), no correlation was indicated between relative maturity and scald percentage from bivariate analysis (Fig. 3). A phenotype value was calculated to account for the linear relationship between scald percentage and relative maturity by determining the deviation from the regression of scald percentage on relative maturity (DRSRRM) and this was used as a phenotype for QTL analysis in our field trials.

### QTLs for relative maturity and plant height

Four QTLs for relative maturity were identified in the Yerong/Franklin population consistently across all three years (Fig. 4 and Table 3): two on chromosome 2H, one on chromosome 5H and another one on chromosome 7H. These four QTLs explained more than 40% of the total phenotypic variance for relative maturity. One QTL for plant height in 2017 was identified on chromosome 3H that explained 14.1% of the total phenotypic variance, with a LOD value of 5.7 (Table 3).

**Fig. 4.**
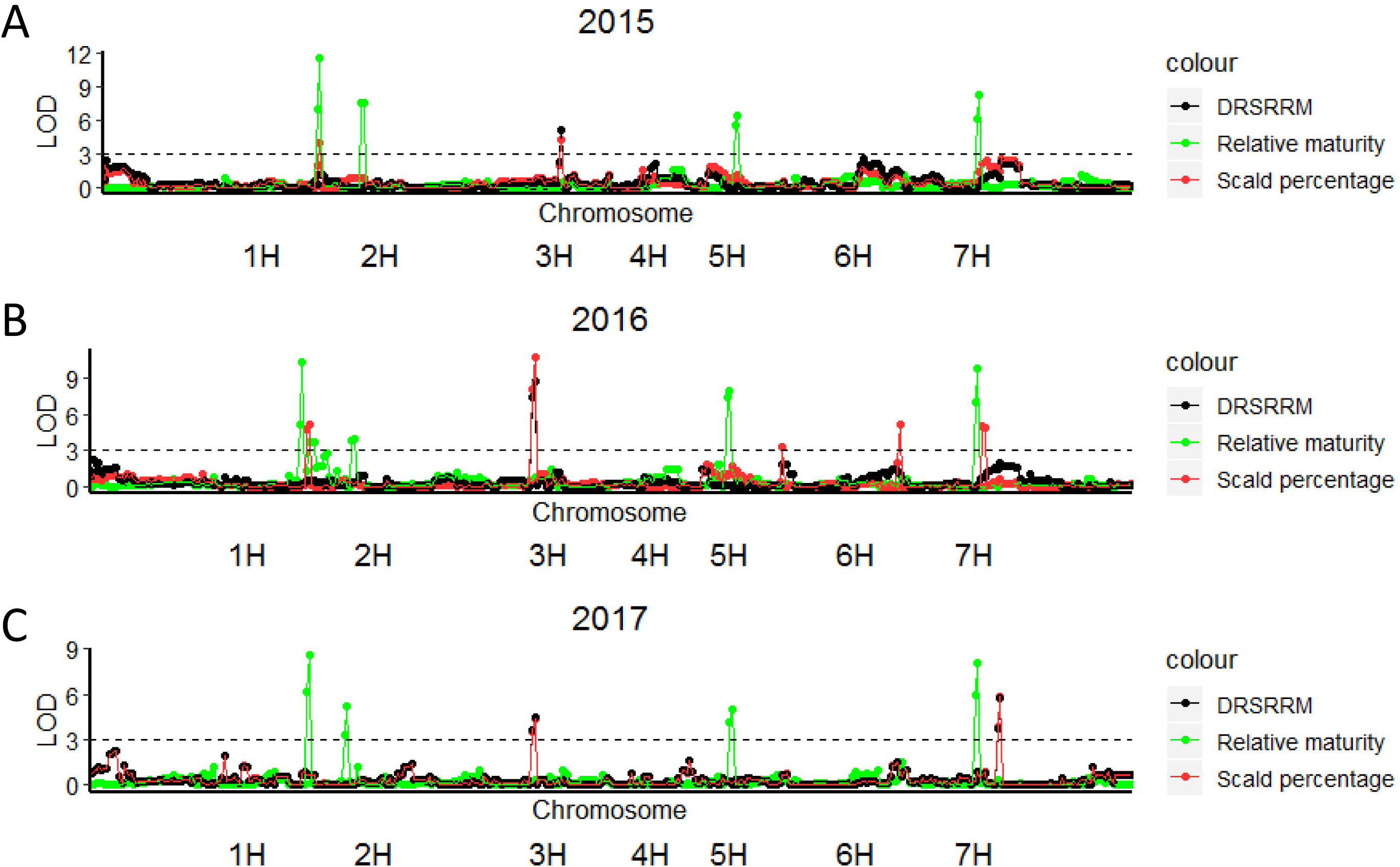
LOD values of QTLs detected for Zadoks score, scald percentage and DRSRRM by MQM mapping across three years field experiments in DH population of Yerong/Franklin. The LOD values of each marker were plotted against the chromosomes

### QTL for scald resistance in the field

In 2015, one QTL for scald percentage on chromosome 2H (QTLSP-2H-2015) was located at the same position as the 2H QTL for relative maturity (QTLZ-2H-2015). In 2016, QTLSP-2H-2016 and QTLZ-2H-2016 were also mapped at the same position on chromosome 2H (Fig. 4 and Table 3). Markers associated with QTLSP-7H-2016 and QTLZ-7H-2016 were also located close to each other (Fig. 4 and Table 3). QTLs for scald percentage were not mapped at the same position as QTLs for relative maturity in 2017.

The QTLs for DRSRRM in 2016 (QTLSR-3H-2016) and 2017 (QTLSR-3H-2017) were located at 178.7 Mb in the same position as QTL-WAIYerong-3H, and the resistant allele was also derived from the Yerong parent. The QTL for DRSRRM (QTL-WAIYerong-3H) explained 20.7% of the phenotypic variance in 2016 (LOD value 8.8; Fig. 4 and Table 3). In 2015, the QTL for DRSRRM on chromosome 3H (QTLSR-3H-2015) was 15 cM away from this major QTL on the linkage map, at 503.4 Mb on the barley physical map. This QTL was still more than 50 Mb away from *Rrs1*, between *Rrs1* (448.4 Mb) and *Rrs4* (523.0 Mb) on chromosome 3H, explaining 14.8% of phenotypic variance. QTLs were identified for DRSRRM on chromosome 3H across the three years at the same location as the QTLs detected for scald percentage. In 2017, a novel QTL for scald resistance (both scald percentage and DRSRRM) was identified on chromosome 7H from Franklin, locating at 69.9 Mb on the barley physical map. This QTL explained 12.9 % phenotypic variance with a LOD value of 5.8.

## Discussion

### Scald resistance at seedling stage

Differential screening conducted in this study showed that the combination of major resistance genes *Rrs1* and *Rrs2* in the variety Atlas46 provides resistance to 10 out of 11 *R. commune* isolates from southern NSW. These results confirmed the effectiveness of pyramiding major resistance genes into one variety in providing broad protection against *R. commune* ^7,33^. However, resistance gene combinations need to be found that do not rapidly select for the corresponding virulence gene combinations in the pathogen population. This remains a challenge with only a limited understanding of the gene for gene interaction between *R. commune* and its host. We have identified specific *R. commune* isolates to distinguish the presence of *Rrs1* or *Rrs2*. While isolates such as WAI2840 posed a threat to reliance on just these two major genes, they are valuable in detecting seedling resistances other than *Rrs1* and *Rrs2* for use in resistance breeding.

Five of the *R. commune* isolates screened in this study were avirulent against Turk and virulent against Yerong, indicating that the *Rrs1* allele from Turk is not present in Yerong. Yerong was resistant to isolate WAI2471, and had moderate levels of resistance against WAI453, WAI1245, WAI2439 and WAI2473. Further QTL analysis located a major QTL for scald resistance against four different isolates at the seedling stage to chromosome 3H, which originated from Yerong. This QTL-WAIYerong-3H explained more than 46% of the phenotypic variation and is located at the same position of a QTL (QSc.YeFr-3H) identified using adult plant data in a previous study from the same DH population in Tasmania, Australia ^23^. However, Li and Zhou (2011) were unable to distinguish this QTL from Yerong from that of *Rrs1*.

The complete reference barley genome sequence enabled the projection of QTLs from different populations onto one barley physical map ^34^. In our study, flanking markers of the QTL-WAIYerong-3H, bPb-0068 and Bmag0006, are mapped at 120.3 and 178.7 Mb on the physical map. The position is some distance away from previously fine mapped *Rrs1* which is located at a position of 448.4 Mb on chromosome 3H (Fig. 2) ^12^. This also suggested the scald resistance QTL-WAIYerong-3H is different from *Rrs1*. Multiple major and minor scald resistance genes or QTLs have been identified on chromosome 3H, close to the *Rrs1* locus ^12,16–20^. Among these QTL, the QTL-IA3H from cultivar Abyssinian was identified in studies of seedling resistance to four *R. commune* isolates ^19^. A QTL Rrs1BC240 for *R. commune* resistance from wild barley *H. spontaneum* was also identified at the position of *Rrs1* ^17^. While ICARDA4 and ICARDA9 showed resistance to all isolates at seedling stage in one study ^7^ and QTLs identified from them were mapped at 383.9 Mb on physical map ^17^, the identity of these resistances remain unknown.

A scald resistance QTL in a similar position to the QTL-WAIYerong-3H (Fig. 2) was published by Coulter, et al. ^20^. QTL qSUK7_3, contributed by the variety Steptoe based on detached leaf assays, was located 3 Mb away from QTL-WAIYerong-3H. The authors suggested this resistance to be different from *Rrs1* based on differential isolate reactions, but were unable to differentiate its map position from that of *Rrs1* ^20^. Further experiments are required to resolve whether the QTL-WAIYerong-3H is the same or different to the gene found in Steptoe on chromosome 3H.

### Disease escape under field conditions

Our study illustrates the potentially confounding effects that relative maturity can have when phenotyping for disease resistance. The observed differences between genotypes may be conferred by both host resistance and mechanisms that lead to disease escape, such as plant maturity, plant height and canopy structure ^8,15^. For example, Zhan, et al. ^8^ postulated that later flowering time and taller plant height can physically slow the upward spread of splash-dispersed *R. commune* and contribute to disease escape. While disease escape traits are important, we sought to identify sources of resistance that are independent of relative maturity, as these QTL are more likely to be useful in breeding programs selecting for a constrained window of flowering time for their target environments.

In this study the positions of all four QTLs for relative maturity were co-located with known phenology QTLs. QTLZ-2H.1 was co-located with a QTL for heading date detected in a TX9425/Franklin population ^35^ and this relative maturity allele was derived from Franklin. This QTL is close to another flowering-time QTL, pseudo-response regulator *Ppd-H1* ^36^, which is located 7 cM away on the consensus map. QTLZ-2H.2 was located at the same position as another QTL for heading date detected by Nduulu, et al. ^37^. QTLZ-5H and QTLZ-7H were located at the same positions as the flowering time genes *eam5* and *Eps-7S*, respectively ^38^.

All of these loci have also been implicated to be associated with resistance at seedling or adult plant stages to five biotrophic or necrotrophic pathogens. von Korff, et al. ^39^ identified QTLs for scald, leaf rust and powdery mildew resistance at a position similar to that of *Ppd-H1*. QTLZ-2H.2 was located at the same position as a QTL for Fusarium head blight resistance detected by Nduulu, et al. ^37^. The results of Nduulu, et al. ^37^ also indicated that the QTLs for heading date and Fusarium head blight resistance were tightly linked rather than pleiotropic. A new gene conferring adult plant resistance to leaf rust, *Rph23*, was identified in the same Yerong/Franklin population used in this study on chromosome 7H, co-located with the QTL QTLZ-7H detected in this study ^40^. Another QTL for stem rust resistance at the seedling stage was also mapped to the same position as QTLZ-7H from Yerong/Franklin population ^41^.

A QTL for plant height was mapped to chromosome 2H at the same position as QTLZ-2H.2 in the Yerong/Franklin population by Xue, et al. ^42^. Interestingly, the QTL for plant height identified in the Yerong/Franklin population is at the same position as a QTL for plant height in a CM72/Gairdner population ^43^. Mature plant height was measured in only one of the experiments in this study (in 2017), and a correlation between mature plant height and scald resistance was not observed (data not shown), and the QTL identified for plant height did not co-locate with scald resistance QTL (Table 3).

### Modelling relative maturity together with disease resistance

The deviation from the regression of infection on important confounding traits has been employed previously as an effective phenotype for disease resistance ^9,31^. Van Beuningen and Kohli ^31^ in particular included linear functions for both heading date and plant height against Septoria tritici blotch (STB) infection in wheat. They aimed to calculate a phenotype that captured components of resistance that did not depend on those two traits, and represented a better approximation of genetic resistance from the experiments in their study. The residuals from a generalized linear model in which STB percentages were fitted to all escape related traits, including heading date, plant height, leaf spacing and leaf morphology, were used as the indicators of disease resistance to analyse the STB resistance in a set of wheat lines ^44^. Chartrain, et al. ^45^ identified a QTL for partial resistance to STB in wheat by using residuals from multiple regression on relative maturity and plant height as their phenotype. QTLs for spot blotch resistance in wheat were detected by using residuals when fitting disease severity as a dependent variable and plant height and days to heading as independent variables in multiple regression to exclude the effects of these traits ^46^. Most of the reported uses of maturity regression residuals phenotype pertain to STB resistance in wheat. This could be due to the well-characterised relationship between STB infection rates and both flowering time and plant height ^47^ and recognition that these confounding traits needed to be accounted for in ascertaining true genetic resistance for STB that would be useful for variety improvement ^48^. In general terms, for phenotypes based on deviation from the regression of a correlated trait (like our DRSRRM measure of resistance) and the approaches described above, the most resistant lines are those with the largest negative values of residuals, also allowing breeders to select resistant lines with desirable relative maturity and plant height ^44,49^. Further, as noted by Van Beuningen and Kohli ^31^, this approach is especially useful where disease resistance is evaluated in experiments at a single date, rather than at critical development stages for each line, especially in genetic material with large variation for confounding traits such as maturity.

In our study, a multiplicative mixed model was used to analyse the field data of scald resistance and relative maturity. A QTL for scald percentage on chromosome 2H were identified at the same position as QTLs for relative maturity; and the major QTL on chromosome 3H was also detected for scald percentage. When DRSRRM was utilised as a trait for QTL analysis, a major QTL was identified on chromosome 3H. This major QTL is co-located with a major QTL detected for scald resistance at seedling stage QTL-WAIYerong-3H. This indicates that by using DRSRRM, QTLs for disease resistance were identified as the confounding effects of relative maturity were removed.

In conclusion, a major QTL QTL-WAIYerong-3H providing scald resistance at both seedling and adult plant growth stages was identified. Results from this study indicate that this QTL-WAIYerong-3H is not *Rrs1*, as differential variety screening indicates the *Rrs1* allele from Turk is not present in Yerong. The flanking markers for this QTL-WAIYerong-3H are located distantly (approx. 270Mb) from the *Rrs1* locus based on the barley physical map. To overcome the confounding effects of relative maturity on adult plant disease resistance, a bivariate approach was used to model the field data of scald resistance and relative maturity. The phenotype derived from the bivariate analysis (DRSRRM) is a more effective trait to detect disease resistance QTLs as it removes the confounding effects of relative maturity. The identified new QTL identified in this study are a useful resource for pyramiding different resistance genes (major or minor) in breeding programs.

## Acknowledgements

The authors thank Tony Goldthorpe, Michael McCaig and Brad Baxter for their expert technical contributions and data collection.

## Author contributions

AM conceived and designed the experiments. MZ, RP and DS provided seeds of the Yerong-Franklin population. XZ performed the phenotype screening. BO, XZ, BAO and AM analysed the data. XZ, BO, BAO, MZ, RP, DS and AM prepared and edited the manuscript.

**Compliance with ethical standards Disclaimer**

## Ethical approval

This article does not contain any studies with human participants or animals performed by the authors.

## References

1 Paulitz, T. & Steffenson, B. J. in Barley: production, improvement and uses (ed S. E. Ullrich) 307–354 (Blackwell Publishing Ltd, 2011).

2 Stefansson, T. S., Serenius, M. & Hallsson, J. H. The genetic diversity of Icelandic populations of two barley leaf pathogens, *Rhynchosporium commune* and *Pyrenophora teres*. European Journal of Plant Pathology 134, 167–180, doi:10.1007/s10658-012-9974-8 (2012).

3 Bouajila, A., Zoghlami, N., Ghorbel, A., Rezgui, S. & Yahyaoui, A. Pathotype and microsatellite analyses reveal new sources of resistance to barley scald in Tunisia. Fems Microbiol Lett 305, 35–41, doi:10.1111/j.1574-6968.2010.01909.x (2010).

4 Goodwin, S. B., Allard, R. W., Hardy, S. A. & Webster, R. K. Hierarchical Structure of Pathogenic Variation among *Rhynchosporium-Secalis* Populations in Idaho and Oregon. Can J Bot 70, 810–817 (1992).

5 Stefansson, T. S., McDonald, B. A. & Willi, Y. The Influence of Genetic Drift and Selection on Quantitative Traits in a Plant Pathogenic Fungus. Plos One 9, doi:10.1371/journal.pone.0112523 (2014).

6 Xi, K. et al. Distribution of pathotypes of *Rhynchosporium secalis* and cultivar reaction on barley in Alberta. Plant Dis 87, 391–396, doi:10.1094/Pdis.2003.87.4.391 (2003).

7 Wallwork, H. & Grcic, M. The use of differential isolates of *Rhynchosporium secalis* to identify resistance to leaf scald in barley. Australasian Plant Pathol. 40, 490–496, doi:10.1007/s13313-011-0065-7 (2011).

8 Zhan, J., Fitt, B. D. L., Pinnschmidt, H. O., Oxley, S. J. P. & Newton, A. C. Resistance, epidemiology and sustainable management of *Rhynchosporium secalis* populations on barley. Plant Pathol 57, 1–14, doi:10.1111/j.1365-3059.2007.01691.x (2008).

9 Brown, J. K. M. Yield penalties of disease resistance in crops. Current Opinion in Plant Biology 5, 339–344, doi:https://doi.org/10.1016/S1369-5266(02)00270-4 (2002).

10 Xi, K., Xue, A. G., Burnett, P. A. & Turkington, T. K. Quantitative resistance of barley cultivars to *Rhynchosporium secalis*. Canadian Journal of Plant Pathology 22, 217–223, doi:10.1080/07060660009500466 (2000).

11 Looseley, M. E. et al. Resistance to *Rhynchosporium commune* in a collection of European spring barley germplasm. Theoretical and Applied Genetics 131, 2513–2528, doi:10.1007/s00122-018-3168-5 (2018).

12 Hofmann, K. et al. Fine mapping of the *Rrs1* resistance locus against scald in two large populations derived from Spanish barley landraces. Theoretical and Applied Genetics 126, 3091–3102, doi:10.1007/s00122-013-2196-4 (2013).

13 Hanemann, A., Schweizer, G. F., Cossu, R., Wicker, T. & Roder, M. S. Fine mapping, physical mapping and development of diagnostic markers for the *Rrs2* scald resistance gene in barley. Theoretical and Applied Genetics 119, 1507–1522, doi:10.1007/s00122-009-1152-9 (2009).

14 Garvin, D. F., Brown, A. H. D. & Burdon, J. J. Inheritance and chromosome locations of scald-resistance genes derived from Iranian and Turkish wild barleys. Theoretical and Applied Genetics 94, 1086–1091, doi:10.1007/s001220050519 (1997).

15 Walters, D. R. et al. Control of foliar diseases in barley: towards an integrated approach. European Journal of Plant Pathology 133, 33–73, doi:http://dx.doi.org/10.1007/s10658-012-9948-x (2012).

16 Bjornstad, A., Patil, V., Tekauz, A., Maroy, A. G. & et al. Resistance to scald (*Rhynchosporium secalis*) in barley (*Hordeum vulgare*) studied by near-isogenic lines: I. markers and differential isolates. Phytopathology 92, 710 (2002).

17 Genger, R. K. et al. Leaf scald resistance genes in *Hordeum vulgare* and *Hordeum vulgare* ssp *spontaneum*: parallels between cultivated and wild barley. Aust J Agr Res 54, 1335–1342, doi:10.1071/ar02230 (2003).

18 Graner, A. & Tekauz, A. RFLP mapping in barley of a dominant gene conferring resistance to scald (*Rhynchosporium secalis*). Theoretical and Applied Genetics 93, 421–425 (1996).

19 Grønnerød, S. et al. Genetic analysis of resistance to barley scald (*Rhynchosporium secalis*) in the Ethiopian line ‘Abyssinian’ (CI668). Euphytica 126, 235–250, doi:10.1023/a:1016368503273 (2002).

20 Coulter, M. et al. Characterisation of barley resistance to rhynchosporium on chromosome 6HS. Theoretical and Applied Genetics 132, 1089–1107, doi:10.1007/s00122-018-3262-8 (2019).

21 Looseley, M. E. et al. Genetic mapping of resistance to Rhynchosporium commune and characterisation of early infection in a winter barley mapping population. Euphytica 203, 337–347, doi:10.1007/s10681-014-1274-2 (2014).

22 Dyck, P. L. & Schaller, C. W. Inheritance of resistance in barley to several physiologic races of the scald fungus. Canadian Journal of Genetics and Cytology 3, 153–164, doi:10.1139/g61-019 (1961).

23 Li, H. B. & Zhou, M. X. Quantitative trait loci controlling barley powdery mildew and scald resistances in two different barley doubled haploid populations. Molecular Breeding 27, 479–490, doi:DOI 10.1007/s11032-010-9445-x (2011).

24 Jackson, L. F. & Webster, R. K. Race differentiation, distribution, and frequency of *Rhynchosporium secalis* in California. Phytopathology 66, 719–7125 (1976).

25 Zadoks, J. C., Chang, T. T. & Konzak, C. F. A decimal code for the growth stages of cereals. Weed research 14, 415–421 (1974).

26 Butler, D., Cullis, B., Gilmour, A. & Gogel, B. ASReml-R reference manual. Queensland Department of Primary Industries and Fisheries: Brisbane, Qld (2009).

27 R: A language and environment for statistical computing (R Foundation for Statistical Computing: Vienna http://www.R-project.org, 2010).

28 Gilmour, A. R., Cullis, B. R. & Verbyla, A. P. Accounting for natural and extraneous variation in the analysis of field experiments. J. Agric. Biol. Environ. Stat 2, 269–293 (1997).

29 Kenward, M. G. & Roger, J. H. Small sample inference for fixed effects from restricted maximum likelihood. Biometrics, 983–997 (1997).

30 Stram, D. O. & Lee, J. W. Variance components testing in the longitudinal mixed effects model. Biometrics, 1171–1177 (1994).

31 Van Beuningen, L. & Kohli, M. Deviation from the regression of infection on heading and height as a measure of resistance to Septoria tritici blotch in wheat. Plant Dis 74, 488–493 (1990).

32 Van Ooijen, J. MapQTL® 6, Software for the mapping of quantitative trait in experiment populations of diploid species. Kyazma BV, Wageningen (2009).

33 Boyd, L. A., Ridout, C., O’Sullivan, D. M., Leach, J. E. & Leung, H. Plant–pathogen interactions: disease resistance in modern agriculture. Trends in Genetics 29, 233–240, doi:http://dx.doi.org/10.1016/j.tig.2012.10.011 (2013).

34 Mascher, M. et al. A chromosome conformation capture ordered sequence of the barley genome. Nature 544, 427–433, doi:10.1038/nature22043 (2017).

35 Wang, J., Yang, J., McNeil, D. L. & Zhou, M. Identification and molecular mapping of a dwarfing gene in barley (*Hordeum vulgare* L.) and its correlation with other agronomic traits. Euphytica 175, 331–342 (2010).

36 Turner, A., Beales, J., Faure, S., Dunford, R. P. & Laurie, D. A. The pseudo-response regulator *Ppd-H1* provides adaptation to photoperiod in barley. Science 310, 1031–1034 (2005).

37 Nduulu, L. M., Mesfin, A., Muehlbauer, G. J. & Smith, K. P. Analysis of the chromosome 2(2H) region of barley associated with the correlated traits Fusarium head blight resistance and heading date. Theoretical and Applied Genetics 115, 561–570, doi:10.1007/s00122-007-0590-5 (2007).

38 Hu, H. et al. Wild barley shows a wider diversity in genes regulating heading date compared with cultivated barley. Euphytica 215, 75, doi:10.1007/s10681-019-2398-1 (2019).

39 von Korff, M., Wang, H., Léon, J. & Pillen, K. AB-QTL analysis in spring barley. I. Detection of resistance genes against powdery mildew, leaf rust and scald introgressed from wild barley. Theoretical and Applied Genetics 111, 583–590, doi:http://dx.doi.org/10.1007/s00122-005-2049-x (2005).

40 Singh, D., Dracatos, P., Derevnina, L., Zhou, M. & Park, R. F. Rph23: A new designated additive adult plant resistance gene to leaf rust in barley on chromosome 7H. Plant Breeding 134, 62–69, doi:10.1111/pbr.12229 (2015).

41 Dracatos, P. M. et al. Inheritance of Prehaustorial Resistance to *Puccinia graminis* f. sp *avenae* in Barley (*Hordeum vulgare* L.). Mol Plant Microbe In 27, 1253–1262, doi:Doi 10.1094/Mpmi-05-14-0140-R (2014).

42 Xue, D.-w. et al. Identification of QTLs for yield and yield components of barley under different growth conditions. Journal of Zhejiang University SCIENCE B 11, 169–176, doi:10.1631/jzus.B0900332 (2010).

43 Xue, D. et al. Identification of QTLs associated with salinity tolerance at late growth stage in barley. Euphytica 169, 187–196, doi:10.1007/s10681-009-9919-2 (2009).

44 Arraiano, L. S. et al. Contributions of disease resistance and escape to the control of septoria tritici blotch of wheat. Plant Pathol 58, 910–922, doi:10.1111/j.1365-3059.2009.02118.x (2009).

45 Chartrain, L., Brading, P. A., Widdowson, J. P. & Brown, J. K. M. Partial Resistance to Septoria Tritici Blotch (*Mycosphaerella graminicola*) in Wheat Cultivars Arina and Riband. Phytopathology 94, 497–504, doi:10.1094/PHYTO.2004.94.5.497 (2004).

46 Singh, V. et al. Phenotyping at hot spots and tagging of QTLs conferring spot blotch resistance in bread wheat. Molecular Biology Reports 43, 1293–1303, doi:10.1007/s11033-016-4066-z (2016).

47 Tavella, C. M. Date of heading and plant height of wheat varieties, as related to septoria leaf blotch damage. Euphytica 27, 577–580, doi:10.1007/bf00043184 (1978).

48 Wilson, R. E. Australian Septoria Nursery 1985 (AUSEN X). 55 (Western Australia Department of Agriculture, Perth, Australia, 1986).

49 Brown, J. K. M. & Rant, J. C. Fitness costs and trade-offs of disease resistance and their consequences for breeding arable crops. Plant Pathol 62, 83–95, doi:10.1111/ppa.12163 (2013).

50 Graner, A., Foroughi-Wehr, B. & Tekauz, A. RFLP mapping of a gene in barley conferring resistance to net blotch (*Pyrenophora teres*). Euphytica 91, 229–234, doi:10.1007/bf00021075 (1996).

51 Patil, V., Bjornstad, A. & Mackey, J. Molecular mapping of a new gene Rrs4(CI11549) for resistance to barley scald (Rhynchosporium secalis). Molecular Breeding 12, 169–183, doi:Doi 10.1023/A:1026076511073 (2003).

